# The *Vibrio* H-ring facilitates the outer membrane penetration of polar-sheathed flagellum

**DOI:** 10.1101/358820

**Authors:** Shiwei Zhu, Tatsuro Nishikino, Seiji Kojima, Michio Homma, Jun Liu

## Abstract

The bacterial flagellum has evolved as one of the most remarkable nanomachines in nature. It provides swimming and swarming motilities that are often essential for the bacterial life cycle and for pathogenesis. Many bacteria such as *Salmonella* and *Vibrio* species use flagella as an external propeller to move to favorable environments, while spirochetes utilize internal periplasmic flagella to drive a serpentine movement of the cell bodies through tissues. Here we use cryo-electron tomography to visualize the polar-sheathed flagellum of *Vibrio alginolyticus* with particular focus on a *Vibrio* specific feature, the H-ring. We characterized the H-ring by identifying its two components FlgT and FlgO. Surprisingly, we discovered that the majority of flagella are located within the periplasmic space in the absence of the H-ring, which are dramatically different from external flagella in wild-type cells. Our results indicate the H-ring has a novel function in facilitating the penetration of the outer membrane and the assembly of the external sheathed flagella. This unexpected finding is however consistent with the notion that the flagella have evolved to adapt highly diverse needs by receiving or removing accessary genes.

**Significance Statement:** Flagellum is the major organelle for motility in many bacterial species. While most bacteria possess external flagella such as the multiple peritrichous flagella found in *Escherichia coli and Salmonella enterica* or the single polar-sheathed flagellum in *Vibrio* spp., spirochetes uniquely assemble periplasmic flagella, which are embedded between their inner and outer membranes. Here, we show for the first time that the external flagella in *Vibrio alginolyticus* can be changed as periplasmic flagella by deleting two flagellar genes. The discovery here may provide a new paradigm to understand the molecular basis underlying flagella assembly, diversity, and evolution.

## Introduction

Flagellum is the major organelle for motility in many bacterial species. It is arguably one of the most complex nanomachines in the bacterial kingdom. Flagella from different species share a conserved core, but also adapt profound variation to accommodate different needs or functions (1, 2). While most bacteria possess external flagella such as the multiple peritrichous flagella found in *Escherichia coli* and *Salmonella enterica* or the single polar-sheathed flagellum in *Vibrio* spp., spirochetes uniquely assemble periplasmic flagella, which are embedded between their inner and outer membranes (3). The flagella have been therefore a paradigm for understanding of evolution and adaptation of bacterial nanomachines (4).

Peritrichous flagella have been extensively studied in *E. coli* and *Salmonella* (5–9). The flagellum is composed of a long helical filament, a hook, and a motor. The motor is a complex macromolecular assembly composed of several ring structures around a rod, which functions as a drive shaft. The MS-ring consists of multiple copies of a single protein FliF and is embedded in the inner membrane. The C-ring is assembled in the cytoplasm and is essential for torque generation and the clockwise/counterclockwise switching of the direction of rotation. A flagellar type-III export apparatus is located underneath the MS- and C-rings. The L-ring is located in the outer membrane. The P-ring is located in the periplasmic space and interacts with the peptidoglycan layer. The P- and L-rings form a bushing at the distal end of the rod. The rotation of the flagella is driven through an interaction between the rotor and the surrounding stator subunits. Two membrane proteins (MotA and MotB) form the stator complex. Powered by the proton motive force, the stator generates the torque required to rotate the motor, the hook, and the filament. The polar-sheathed flagellum from *Vibrio* species is quite different from the peritrichous flagella in *E. coli* and *Salmonella* (9–11). The polar-sheathed flagellum utilizes a sodium ion gradient as the energy resource for rotation and exhibits a remarkably fast speeds of up to 1700 Hz (12). Compared to the flagella in *E. coli* and *Salmonella*, the polar-sheathed flagellum in *Vibrio* spp. possesses extra ring-like structures known as the T-ring and the H-ring (13, 14). They are essential for high speed rotation of the *Vibrio* flagella (15). The T-ring is located right next to the P-ring and is important for incorporating the sodium-driven stator units into the basal body (13). The H-ring is known to be adjacent to the L-ring. FlgT was the first protein identified to be involved in the formation of the H-ring (15). However, the exact structure and function of the H-ring remained to be defined in detail.

Spirochetes are a group of bacteria with distinctive morphology and motility (3). The motility of the spirochetes is driven by periplasmic flagella, which are enclosed between the inner and outer membranes. The unique location clearly distinguishes the periplasmic flagella from the external flagella in *E. coli* and *Vibrio*. Interestingly, the highly conserved flagellar type III secretion system has been utilized to assemble the rod, the hook, and the filament in both periplasmic flagella and external flagella (3, 16, 17). Therefore, one of the main differences between two flagellar systems is whether or not the flagella penetrate the outer membrane. It is of great interest to identify genes involved in outer membrane penetration.

*Vibrio alginolyticus* is a great model system to study polar-sheathed flagella (10, 18). In particular, cryo-electron tomography (cryo-ET) was recently utilized to visualize the *in situ* structure of sheathed flagella in *V. alginolyticus* that revealed distinct *Vibrio* specific features: the membrane sheath, the O-ring, the T-ring and the H-ring (11). Our previous studies provided structural evidence that MotX and MotY form the T-ring adjacent to the P-ring (11, 13). Here we attempt to understand structure and function of the H-ring by systemically characterizing two mutants lacking *flgT* or *flgO*, respectively. To our surprise, we found that periplasmic flagella assemble in both mutants. The unexpected observation suggests that the H-ring is essential for outer membrane penetration and assembly of the polar flagellum in *Vibrio*. More importantly, the new findings provide a basis for the further understanding of flagellar assembly and evolution.

## Results

### FlgO and FlgT are involved in the H-ring formation

The H-ring is a *Vibrio*-specific flagellar feature that is important for motility. Our recent studies of the *V. alginolyticus* flagellar motor showed that the H-ring is a large disk underneath the outer membrane (11). FlgT is the first protein known to be involved in H-ring formation (14, 15). FlgT is a small protein that might be limited to the proximal part of the H-ring. FlgO and FlgP are two outer membrane lipoproteins required for flagellum stability and motility of *V. cholerae*, as the *flgO* and *flgP* mutants have reduced motility and fewer external flagella (19). The averaged motor structure of *Vibrio fischeri* Δ*flgP* mutant showed that the PL-ring, together with the T-ring, were visible (Morgan et al., 2016). We therefore hypothesize that both FlgO and FlgP might be involved in the formation of the H-ring complex in *Vibrio*. We constructed Δ*flgO* and Δ*flg*T mutant in the background of the multi-polar flagellated strain, respectively (Table 1). The Δ*flgO* mutant cells are less motile in a soft agar plate while expression of a His-tagged *flgO*+ allele complemented the Δ*flgO* allele for motility and expression (Fig. 1A) as detected by western blot using anti-his tag antibody (Fig. 1B).

**Table 1.** Bacterial strains and plasmids 369 used in this study

| Strain or plasmid | Genotype or description | Reference or source |
| --- | --- | --- |
| <u>*V. alginolyticus*</u> |  |  |
| VIO5 | VIK4 (Rif<sup>r</sup> Pof<sup>+</sup> Laf<sup>-</sup>) | (37) |
| KK148 | VIO5 *flhG* (multi-Pof<sup>+</sup>) | (38) |
| TH7 | KK148 Δ*flgT* | (14) |
| NMB337 | KK148 Δ*flgO* | This study |
| <u>*E. coli*</u> |  |  |
| DH5α | Recipient for DNA manipulation | |
| β3914 | Recipient for conjugational transfer of pSW7848 | (39) |
| <u>Plasmids</u> |  | Promega |
| pGEM-T Easy | Cloning vector, Amp<sup>r</sup> | (40) |
| pSW7848 | Suicide vector, (oriVR6Kγ oriTRP4 *araC*-P<sub>BAD</sub>-*ccdB*), Cm<sup>r</sup> | This study |
| pBAD33 | Cm<sup>r</sup>, P<sub>BAD</sub> | This study |
| pTSK127 | pGEM-T Easy-Δ*flgO* | (18) |
| pTSK127_2 | pSW7848-Δ*flgO* | This study |
| pTY57 | Cm<sup>r</sup>, P<sub>BAD</sub> with a multicloning site of pBAD24 |  |
| pTSK128 | pTY57-*flgO*::His<sub>6</sub> |  |

A: Rif<sup>r</sup>, rifampin resistant; Pof<sup>+</sup>, normal polar flagellar formation; Laf<sup>-</sup>, defective in lateral flagellar formation; multi-Pof<sup>+</sup>, multiple polar flagellar formation. Amp<sup>r</sup>, ampicillin resistant; Cm<sup>r</sup>, chloramphenicol resistant; P<sub>BAD</sub>, arabinose promoter.

**Figure 1.**
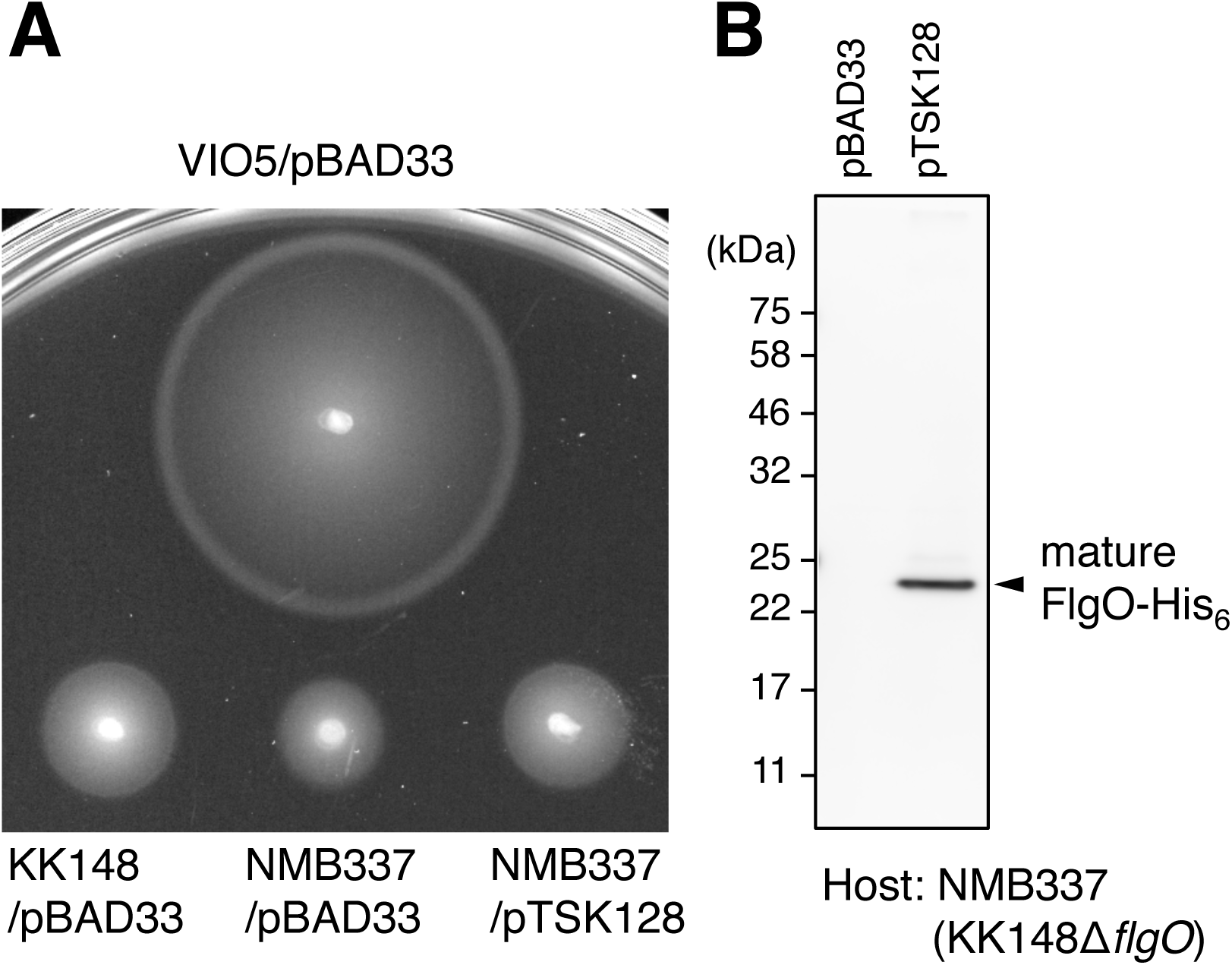
Lack of FlgO results in reduced motility. **(A)** Motility of cells in soft agar. Two μl aliquots of overnight cultures of each strain were spotted onto 0.25% soft agar VPG plate containing 2.5 μg/ml chloramphenicol and 0.02% (wt/vol) LL-arabinose, and the plate was incubated at 30°C for 7 hours. Deletion of *flgO* from the strain KK148 resulted in reduced motility, and ectopic expression of FlgO fused with hexa-histidine tag at the C-terminus (FlgO-His6) from the arabinose-inducible plasmid pTSK128 restored motility (protein expression was confirmed in (B)). VIO5 is the wild type strain for polar flagellar motility; KK148 is multipolar flagellar strain, and the parent of NMB337. The plasmid pBAD33 was used as the empty vector control. **(B)** Immunoblot analysis. Whole cell lysates were separated by SDS-PAGE and transferred onto the PVDF membrane, and his-tagged proteins were detected by anti-His tag antibody. The FlgO-His6 protein was detected at the size equivalent to its mature form (indicated as the filled arrow). Experiments were conducted 3 times and the typical results are shown here.

To decipher whether deletion of *flgO* affects the assembly of the polar-sheathed flagellum and the formation of the H-ring, we examined Δ*flgO* mutant cells by cryo-ET. Polar-sheathed flagella are clearly visible in the Δ*flgO* mutant (Fig. 2, Table 1). We identified flagellar motor structures from tomograms and determined the *in situ* motor structure from the Δ*flgO* strain using sub-tomogram averaging (Fig. 2). Compared to the motor structure from wild type, the distal part of the H-ring density is absent in the Δ*flgO* motor (Fig. 2G, H, J, K and L). Thus, our data suggest that FlgO is the protein component that is primarily responsible for the distal part of the H-ring. Furthermore, the distal part of the H-ring seems to anchor the whole disk onto the inner leaflet of the outer membrane, because the smaller H-ring in the Δ*flgO* motor appears to less tightly associate with the outer membrane (Fig. 2G, H).

**Figure 2.**
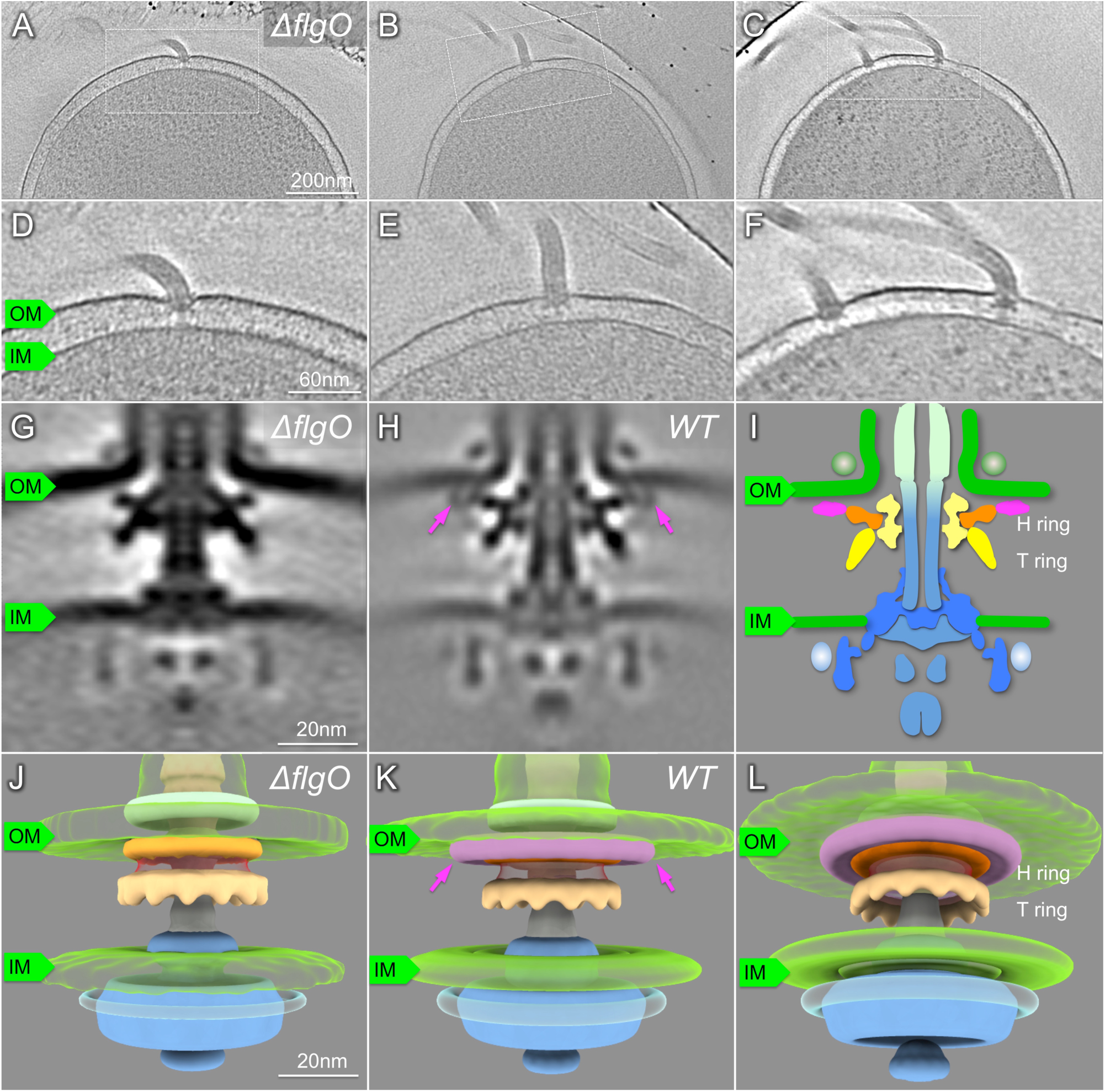
Characterization of the Δ*flgO* flagellum *in situ* by cryo-ET. (A-C) A representative slice of a 3D reconstruction of the *V. alginolyticus* Δ*flgO* strain KK148 with multiple polar flagella. (D-F) Zoom-in views of the slices are shown in A-C. (G) A slice of a sub-tomogram average of the flagellar motor. (H) A slice of a sub-tomogram average of the flagellar motor in KK148. The structural difference between panels G and H is indicated with purple arrows (I) Schematic model of the *Vibrio* motor. (J) 3D surface renderings of (G). (K, L) 3D surface renderings of (H). The H-ring is labeled in orange and pink colors, separately; The T-ring is colored yellow; OM, outer membrane; IM, inner membrane.

To further understand the role of the H-ring on flagellar formation, we visualized a Δ*flgT* mutant using cryo-ET, as FlgT is involved in the formation of the H-ring (14). Indeed, the entire H-ring density is absent in the tomograms of the *flgT* mutant cells, while T-ring density remains visible (Fig. 3). This is consistent with the previous observation that the H-ring is not visible in a purified basal body of the Δ*flgT* mutant by negative stain electron microscopy (14). Together with the results from Δ*flgO* cells, we determined that FlgO is responsible for the distal part of the H-ring and FlgT is essential for the proximal part of the H-ring. Thus, FlgT and FlgO together with FlgP contribute to the formation of the H-ring.

**Figure 3.**
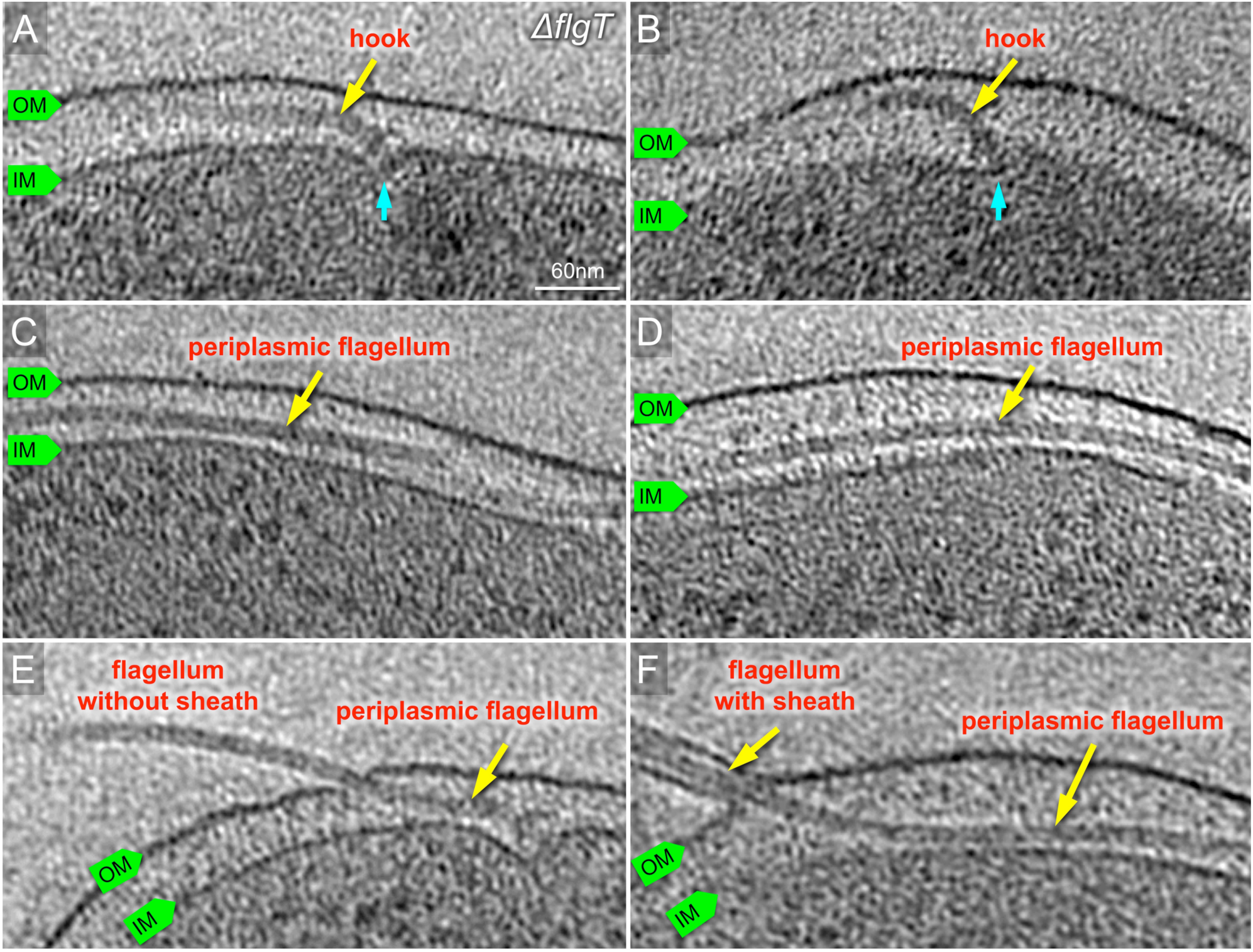
Characterization of the Δ*flgT* flagellum in situ by cryo-ET. (A, B) Representative slices of tomograms from KK148 Δ*flgT* cells. The motor is visible beneath of outer membrane. The motor is colored in cyan and the hook in yellow. (C, D) Representative slices of from KK148 Δ*flgT* cells. The flagellar filament is visible in the periplasmic space and labeled in red. (E) The flagellar filament is extended in the periplasmic space and penetrates the outer membrane without a sheath. (F) The flagellar filament penetrates from the periplasm and sheathed.

### The H-ring plays an essential role in flagella assembly and bacterial motility

The H-ring is tightly associated with the outer membrane (Fig. 2G and 2H). It has been suggested that the H-ring is important for torque generation and bacterial motility (20), although the exact role of the H-ring remains to be determined. To understand the function of the H-ring in more detail, we thoroughly screened tomograms from the Δ*flgO* mutant cells. Surprisingly, most Δ*flgO* mutant cells possess polar-sheathed flagella as those from wild type. However, about 10% of the Δ*flgO* mutant cells displayed both polar-sheathed flagella and periplasmic flagella (Fig. S1). This observation suggested that the H-ring was likely involved in flagellar assembly, especially in penetration of the outer membrane to enable the formation of the external-sheathed flagella.

To further understand the relationship of the H-ring and flagellar assembly, we carefully examined over several hundred reconstructions from Δ*flgT* mutant cells. We found that many hooks are severely bent beneath the PG layer and many filaments are located in the periplasmic space (Fig. S2 and Fig. 3). Since some of the filaments are much longer than the cell body, they often protrude through the PG layer and the outer membrane at the region far from the basal body (Fig. 3). Less external flagella were found on the Δ*flgT* cells than on wild type cells. In total, ~80% of 354 flagella found in the Δ*flgT* cells were located in periplasmic space. Compared to ~10% periplasmic flagella in the Δ*flgO* cells and none in wild type, lack of the H-ring had a profound impact on flagellar assembly and location. Although this result was not previously visualized, it is consistent with the early observation that flagellated cells were rare in the *flgT* mutant cells (21, 22).

### Whole-cell reconstructions show different flagellar assembly and location in wild type and Δ*flgT* mutant cells

For a comprehensive understanding of the impact of the H-ring, we generated the whole-cell reconstructions from the Δ*flgT* mutant and wild type cells. Two flagella are found in the periplasmic region of the Δ*flgT* cells (Fig. 4A-D). One flagellar filament extended into the cell wall and extruded through the outer membrane and was covered by the sheath at the cell pole (Fig. 4A-D). Another flagellar filament folded back towards the cell body and stayed in the periplasmic space. In contrast, no periplasmic flagella were visible in wild type (Fig. 4E, F). Three flagella directly assembled at the pole and penetrated across the outer membrane to form the long external filaments covered with the outer membrane sheath (Fig. 4E, F). Together, we conclude that the loss of the H-ring has a substantial impact on the assembly of the polar-sheathed flagella.

**Figure 4.**
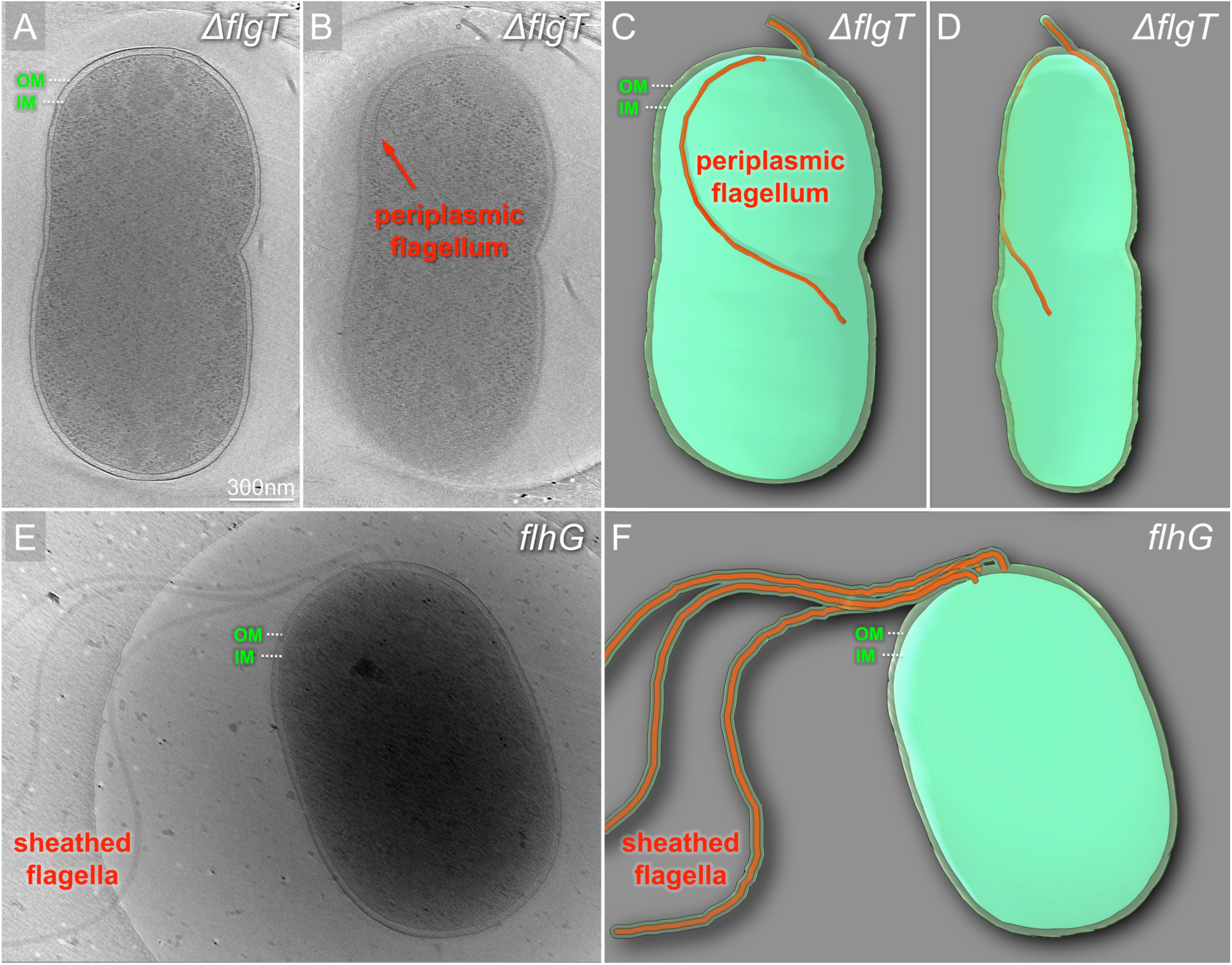
Cryo-ET reconstructions of the whole cells from KK148 and KK148 Δ*flgT* exhibiting dramatic differences in flagella structures. (A, B) Tomographic slice of KK148 Δ*flgT* shown in different layers of the tomogram. (C, D) A and B in a 3D segmentation to show the periplasmic flagella. (E) A representative tomogram slice of a KK148 whole cell shows multiple polar-sheathed flagella. (F) A 3D segmentation of the (E). OM, outer membrane; IM, inner membrane.

## Discussion

The flagella have evolved as the main organelles for motility in many bacteria. Recent studies based on genome sequences and *in situ* structural studies by cryo-ET have demonstrated that while the flagella possess a conserved core, the overall flagellar structures appear to be strikingly diverse in different bacterial species (1). For example, a large cage-like structure surrounds the P- /L- rings in the *H. pylori* flagella (23). *Vibrio* flagella possess the unique H-/T-rings essential for its motility (11, 20). In spirochetes, a large collar-like periplasmic structure is necessary for the assembly of the periplasmic flagella and the unique spirochete motility (24–26). The distinction between periplasmic flagella in spirochetes and external flagella in other species is among the most noticeable differences among different bacterial flagella. To better understand the structure, function and evolution of the flagellum, it is of particular interest to uncover unique aspects of flagella in different bacterial species. Using *V. alginolyticus* as a model system, we previously analyzed the *Vibrio* specific T-ring, which is vital for higher torque generation and faster motility in *Vibrio*. Here we revealed that the *Vibrio* specific H-ring is required for flagellar morphogenesis and assembly.

### Novel structure and function of the H-ring *in Vibrio*

Our studies clearly indicated that FlgO forms the distal part of the H-ring and FlgT is responsible for the proximal part. Another protein component of the H-ring might be FlgP, as it is a lipoprotein localized to the outer membrane in *V. cholera* (19, 27). Recent cryo-ET studies of a Δ*flgP* mutant from *Vibrio fischeri* provided evidence that most of the H-ring is absent (20), while the small density adjacent to the L-ring remains. Together with our results from Δ*flgO* and Δ*flgT*, it is very likely that FlgT, FlgP, and FlgO are directly involved in the proximal, middle, and distal parts of the H-ring, respectively.

The H-ring has been suggested to be important for high torque generation (20). The H-ring is not only visible in *Vibrio* species, but also present in *Aeromonas hydrophila* species (28). Since flagella in both species are sodium driven, which are known to generate greater torque than flagellar motors driven by proton flow (29). The sodium-driven flagella also evolved additional accessary structures such as the T-ring and the H-ring to support higher torque generation (30). However, we found that reduced motility due to FlgT or FlgO dysfunction is attributed to a significant change on flagellar morphogenesis from polar flagella to periplasmic flagella in addition to an effect on torque generation. Thus, the H-ring plays important roles not only in stabilizing flagellar motors on the outer membrane but also in facilitating outer membrane penetration by extracellular flagella.

### Outer membrane penetration and implication on flagella evolution

The observation that both Δ*flgT and* Δ*flgO* mutant cells grew periplasmic flagella was surprising. Although it has been reported previously that conversion from external to periplasmic flagella could result from single amino acid substitutions in the flagellar rod protein FlgG (31), our observation is quite different. FlgG is a conserved core protein and the length of flagellar rod composed of multiple FlgG is relatively constant in both spirochete periplasmic flagella and external flagella (32). On the contrary, both FlgT and FlgO are not conserved core proteins (33). In the Δ*flgO* mutant cells, only the distal portion of the H-ring structure is absent. The overall structure of the flagellar motor is similar to that in wild type and most flagella are able to penetrate the outer membrane and form the external flagella covered by the sheath. However, 10% of flagella fail to penetrate the outer membrane resulting in the formation of periplasmic flagella. In the absence of the entire H-ring, as was observed in a Δ*flgT* mutant strain, the majority of flagella fail to penetrate the outer membrane and form the normal, sheathed flagella. Instead, they form periplasmic flagella. Some of them are much longer than the cell body and appear to violently protrude from the cell wall without any bushing such as the PL-rings. Thus, the H-ring plays important roles not only in stabilizing flagellar motors on the outer membrane but also in facilitating outer membrane penetration by extracellular flagella.

The H-ring is only a part of the large outer membrane complex, which is often structurally variable in different Gram-negative bacterial species and is also absent in both Gram-positive bacteria and spirochetes. The fact that it is possible to generate periplasmic flagella from external flagella by altering the outer membrane complex is particularly interesting. This allows us to speculate that periplasmic flagella might be evolved from extracellular flagella by losing genes involving in the formation of the outer membrane complex. On the other hand, it also raises the possibility that extracellular flagella could evolve from periplasmic flagella by receiving genes critical for the outer membrane complex formation. Thus, the function of outer membrane complex determines the bacterial flagellar morphogenesis: external or periplasmic. During flagellar assembly, the flagellar rod is also playing an important role in penetrating the outer membrane (31, 34). Thus, the change of flagellar morphogenesis would be attributed to the cross-talk between the flagellar rod and the outer membrane complex. How the flagellar rod senses and coordinates with outer membrane complex is await to be revealed.

In summary, we characterized the *Vibrio*-specific H-ring by using cryo-ET and genetic mutations and provided evidence that at least two proteins (FlgO and FlgT) are directly involved in the formation of the H-ring (Fig. 5). Furthermore, we discovered that the H-ring plays a novel function in facilitating the penetration of the outer membrane in *Vibrio* species. Thus, we concluded that the outer membrane complex is not only working as the bushing, but also functioning to an adaptor to flagellar rod to determine the flagellar morphogenesis (Fig. 5). Periplasmic flagella assembled are observed for the first time *in situ*. The discovery here may provide a new paradigm to understand the molecular basis underlying flagella assembly, diversity, and evolution.

**Figure 5.**
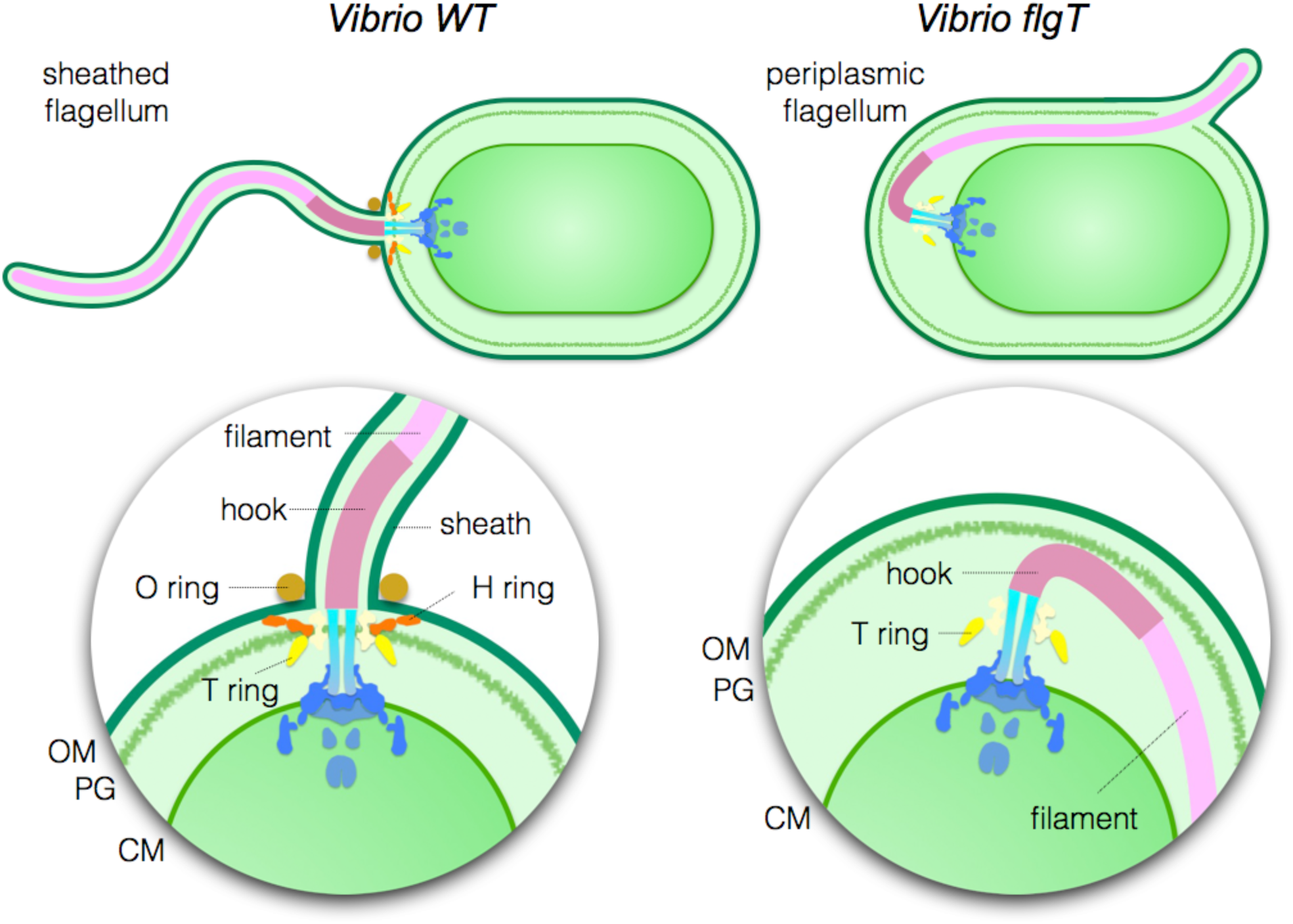
Model of polar-sheathed flagellar assembly. *Vibrio* species have a single polar-sheathed flagellum. The H-ring labeled in red is required in the assembly of the polar-sheathed flagellum, in addition to flagellar stabilization. The dysfunction of FlgT causes the loss of the H-ring and consequently, change the polar flagellum to a periplasmic flagellum. OM, outer membrane; PG, peptidoglycan layer; IM, inner membrane.

## Materials And Methods

### Bacterial Strains, plasmids and growth condition

Bacterial strains used in this study are listed in Table 1. *V. alginolyticus* strains were cultured at 30°C on VC medium [0.5 % (wt/vol) polypeptone, 0.5% (wt/vol) yeast extract, 3% (wt/vol) NaCl, 0.4% (wt/vol) K2HPO4, 0.2% (wt/vol) glucose] or VPG medium [1% (wt/vol) polypeptone, 3% (wt/vol) NaCl, 0.4% (wt/vol) K2HPO4, 0.5% (wt/vol) glycerol]. If needed, chloramphenicol and Larabinose were added at final concentrations of 2.5 μg/mL and 0.02% (wt/vol), respectively. *E. coli* was cultured at 37°C in LB medium [1% (wt/vol) bactotryptone, 0.5% (wt/vol) yeast extract, 0.5% (wt/vol) NaCl]. If needed, chloramphenicol and ampicillin were added at final concentrations of 25 μg/ml and 100 μg/ml, respectively. Introduction of plasmids into *Vibrio* strains were conducted by electroporation as described previously (Kawagishi et al, 1994).

### Construction of the *flgO* deletion strain

The *flgO* deletion strain NMB337 was generated from multi-polar flagellated strain KK148 by homologous recombination with the Δ*flgO* sequence (1,000 bp), which is composed of 500 bp upstream sequence of *flgO* fused with 500 bp downstream sequence of *flgO*, by using the method described previously (35). The Δ*flgO* DNA fragment was amplified by two-step PCR: for upstream sequence using a sense primer 1 (5’- GGGAGCTCATGGATAAATATCGACGCGAA-3’) containing a *Sac*I site and an antisense primer 2 (5’-CATGCTTCTATCGGTTTGATTCTCCAGATAATC-3’), and for downstream sequence using a sense primer 3 (5’-GAGAATCAAACCGATAGAAGCATGAAGAAGTT-3’) and an antisense primer 4 (5’-AAGAGCTCTGTTGCCAATCAGCCG-3’) containing a *Sac*I site. Amplified PCR fragments for upstream and downstream sequence were gel-purified and mixed, then Δ*flgO* DNA fragment was PCR amplified by using a sense primer 1 and an antisense primer 4. The Δ*flgO* fragment was cloned into pGEM-T Easy vector using *Sac*I site to generate pTSK127, and then it was transferred to pSW7848 to generate pTSK127_2. By using the conjugational transfer, pTSK127_2 was introduced into KK148, and Δ*flgO* strains were obtained as described previously (35). The deletion was confirmed by colony PCR and DNA sequencing.

### Motility assay

Two μL of overnight cultures of *V. alginolyticus* cells containing plasmids at 30°C in VC medium with chloramphenicol were spotted on the VPG soft agar plate (VPG medium containing 0.25% [vt/vol] Bact agar with 0.02% (vt/vol) L-arabinose and 2.5μg/ml chloramphenicol). The plate was incubated at 30 °C for 7 hours.

### Detection of proteins by immunoblotting

*Vibrio* cells grown overnight at 30°C in VC medium were re-inoculated at a 100-fold dilution into fresh VPG medium containing 0.02% [vt/vol] Larabinose and 2.5 μg/ml chloramphenicol. Cells were cultured at 30°C for about 3.5 hours, harvested, suspended to an optical density at 660 nm equivalent to 10 in SDS loading buffer and boiled at 95°C for 5 min. These whole cell lysate samples were separated by SDS-PAGE and transferred to polyvinylidene difluoride (PVDF) membrane, and immunoblotting was performed using polyclonal anti-His tag antibody (Medical and Biological Laboratories Co., Ltd., Nagoya Japan).

### Sample preparation for cryo-ET observation

*V. alginolyticus* strains were cultured overnight at 30°C on VC medium and diluted 100× with fresh VC medium and cultured at 30°C at 220 rpm (Taitec, BioShaker BR-23FH). After 5 h, cells were collected and washed 2× and finally diluted with TMN500 medium (50 mM Tris-HCl at pH 7.5, 5 mM glucose, 5 mM MgCl, and 500 mM NaCl). Colloidal gold solution (10 nm diameter) was added to the diluted Vibrio samples to yield a 10× dilution and then deposited on a freshly glow-discharged, holey carbon grid for 1 min. The grid was blotted with filter paper and rapidly plunge-frozen in liquid ethane in a homemade plunger apparatus, as described previously (11).

### Cryo-ET data collection and image processing

The frozen-hydrated specimens of KK148 and TH7 were transferred to a Polara G2 electron microscope and the samples of NMB337 was transfer to Titan Krios electron microscope (FEI). Both microscopes are equipped with a 300-kV field emission gun and a Direct Electron Detector (Gatan K2 Summit). Images collected by Polara G2 electron microscope were observed at 9,000× magnification and at ∼8 μm defocus, resulting in 0.42 nm/pixel. The images taken by Titan Krios electron microscope were collected at a defocus near to 0 um using Volta Phase Plate and the energy filter with 20 eV slit. The data was acquired automatically with SerialEM software (36). During the data collected, when phase shift is out of the range of pi/3~pi2/3, next spot of phase plate will be switched to be charged for use. A total dose of 50 e−/Å2 is distributed among 35 tilt images covering angles from −51° to +51° at tilt steps of 3°. For every single tilt series collection, the dose-fractionated mode was used to generate 8–10 frames per projection image. Collected dose-fractionated data were first subjected to the motion correction program to generate drift-corrected stack files (Li *et al.*, 2013; Morado *et al.*, 2016; Zheng *et al.*, 2017). The stack files were aligned using gold fiducial markers and volumes reconstructed by the weighted back-projection method, using IMOD and Tomo3d software to generate tomograms (Kremer *et al.*, 1996; Agulleiro and Fernandez, 2015). In total, 137 tomograms of TH7 and 114 tomograms of NMB337 were generated.

### Sub-tomogram analysis with i3 package

Bacterial flagellar motors were detected manually, using the i3 program (Winkler, 2007; Winkler *et al.*, 2009). We selected two points on each motor: one point at the C-ring region and another near the flagellar hook. The orientation and geographic coordinates of selected particles were then estimated. In total, 668 sub tomograms of *Vibrio* motors from NMB337 were used to sub-tomogram analysis. The i3 tomographic package was used on the basis of the “alignment by classification” method with missing wedge compensation for generating the averaged structure of the motor, as described previously (11).

### 3D visualization

Tomographic reconstructions were visualized using IMOD (Kremer *et al.*, 1996). UCSF Chimera software was used for 3D surface rendering of subtomogram averages and molecular modeling (Pettersen *et al.*, 2004).

## Acknowledgements

We thank Kelly Hughes for critically reading the manuscript prior to submission. This work was supported by GM107629 from the National Institute of General Medicine, and AU-1714 from the Welch Foundation (to J. L.).

---

**Fig. S1.**
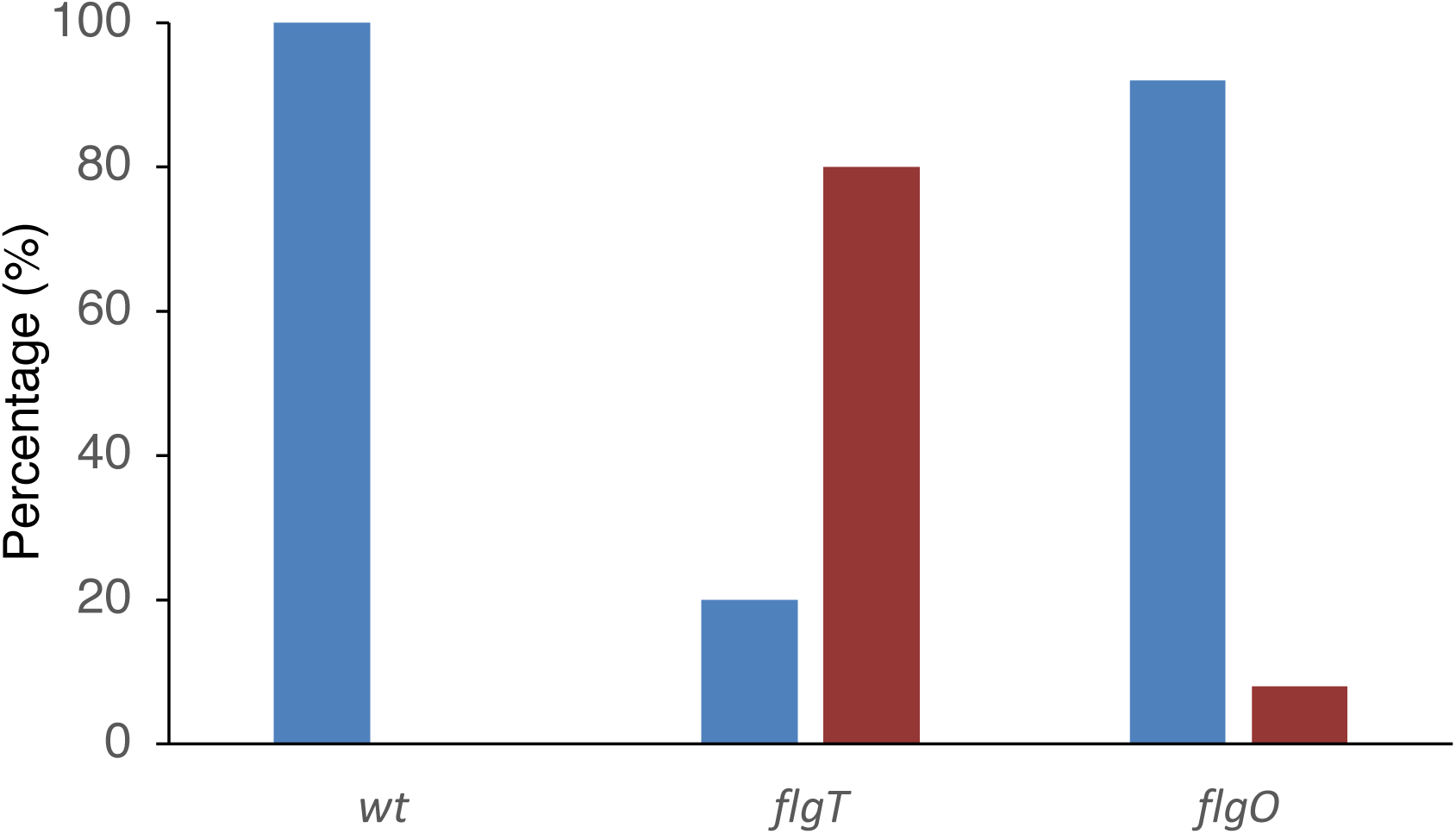
Percentage of periplasmic flagella (red) and polar flagella (blue) found in wild type, Δ*flgT* and Δ*flgO* cells.

**Fig. S2.**
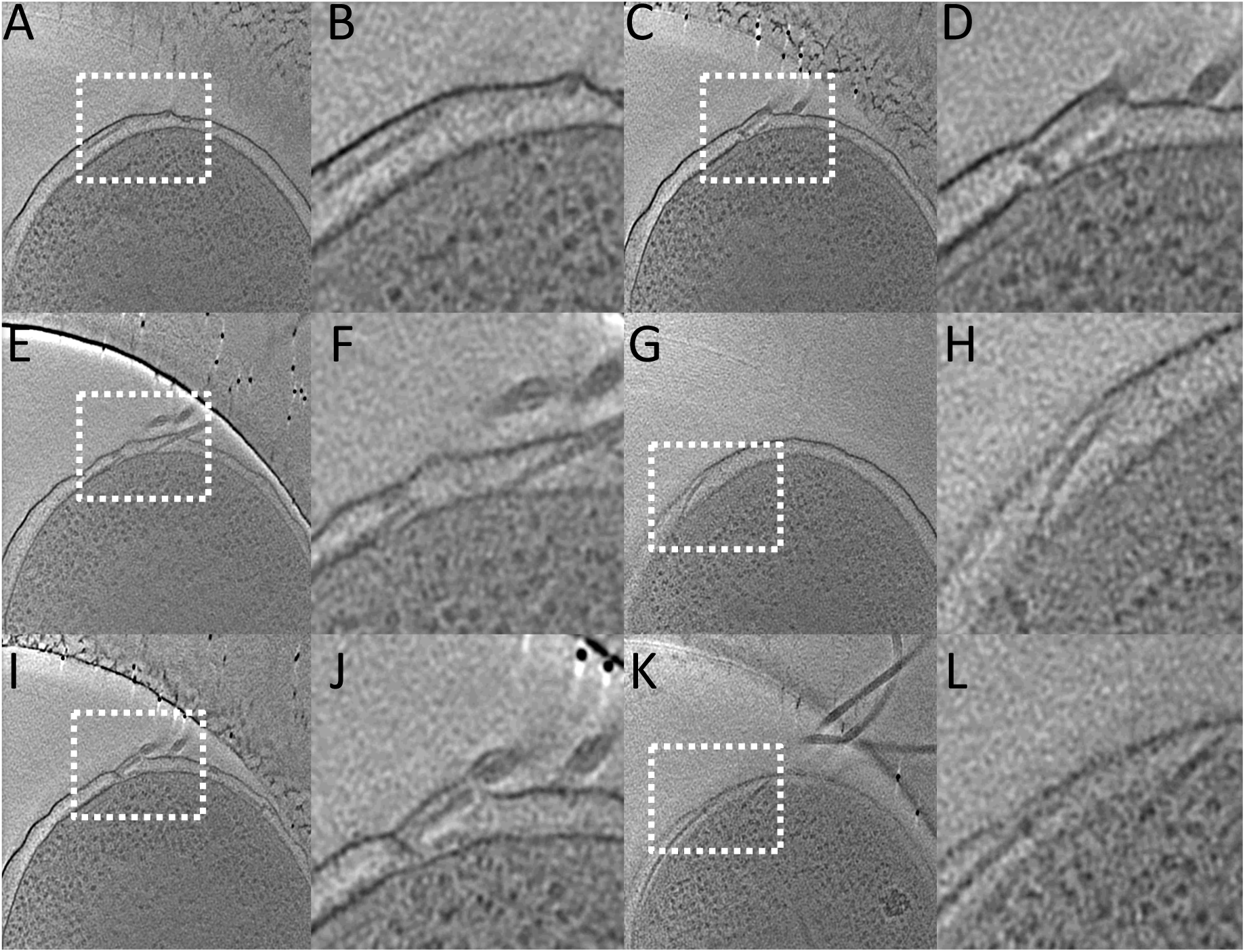
Gallery of periplasmic flagella found in the Δ*flgO* cells.

